# Focused Ultrasound Modulates Dopamine in a Mesolimbic Reward Circuit

**DOI:** 10.1101/2024.02.13.580202

**Authors:** Greatness O. Olaitan, Mallikarjunarao Ganesana, Andrew Strohman, Wendy J. Lynch, Wynn Legon, B. Jill Venton

## Abstract

Dopamine is a neurotransmitter that plays a significant role in reward and motivation. Dysfunction in the mesolimbic dopamine pathway has been linked to a variety of psychiatric disorders, including addiction. Low-intensity focused ultrasound (LIFU) has demonstrated effects on brain activity, but how LIFU affects dopamine neurotransmission is not known. Here, we applied three different intensities (6.5, 13, and 26 W/cm^2^ I_sppa_) of 2-minute LIFU to the prelimbic region (PLC) and measured dopamine in the nucleus accumbens (NAc) core using fast-scan cyclic voltammetry. Two minutes of LIFU sonication at 13 W/cm^2^ to the PLC significantly reduced dopamine release by ∼ 50% for up to 2 hours. However, double the intensity (26 W/cm^2^) resulted in less inhibition (∼30%), and half the intensity (6.5 W/cm^2^) did not result in any inhibition of dopamine. Anatomical controls applying LIFU to the primary somatosensory cortex did not change NAc core dopamine, and applying LIFU to the PLC did not affect dopamine release in the caudate or NAc shell. Histological evaluations showed no evidence of cell damage or death. Modeling of temperature rise demonstrates a maximum temperature change of 0.5°C with 13 W/cm^2^, suggesting that modulation is not due to thermal mechanisms. These studies show that LIFU at a moderate intensity provides a noninvasive, high spatial resolution means to modulate specific mesolimbic circuits that could be used in future studies to target and repair pathways that are dysfunctional in addiction and other psychiatric diseases.

## Introduction

Dopamine is a neuromodulator with many essential roles in the central nervous system.^1,2^ In the mesolimbic system, dopaminergic neurons from the ventral tegmental area (VTA) innervate various brain regions, including the prefrontal cortex (PFC) and nucleus accumbens (NAc), and play a pivotal role in regulating feelings of well-being and reward. ^3–6^ A particularly notable circuitry within the reward pathway is the connection between glutamatergic neurons from the prelimbic region (PLC) of the medial PFC (mPFC) and dopaminergic neurons in the NAc core.^7–9^ This circuitry regulates reward-related motivation by controlling dopamine release in the NAc core. Dysregulation of this specific circuit is observed in reward-related diseases such as addiction and pathological gambling.^10^ Consequently, re-regulating this glutamate-dopamine circuitry could provide a strategic approach to treating neuropsychological diseases involving the mesolimbic dopamine system.

Targeted neuromodulation represents a new frontier in treating neuropsychological diseases, offering precise control over specific neural circuits that traditional pharmacological approaches fail to provide.^11–13^ This specificity is crucial, especially in conditions associated with disruptions in dopamine, considering the diverse roles it plays in reward processing, motor control, and emotional responses.^14^ Noninvasive methods to regulate neural activity, such as transcranial magnetic stimulation (TMS) and transcranial electrical stimulation (TES), have emerged as promising alternatives.^15,16^ Although TMS and TES can exert both excitatory and inhibitory effects on neurotransmission, with some clinical successes, they are limited by poor spatial resolution and an inherent depth-focality tradeoff.^17^ This limitation constrains their ability to target deeper brain areas with precision. In contrast, Low-Intensity Focused Ultrasound (LIFU) utilizes acoustic waves to modulate neuronal activity by focusing these waves on small, specific regions deep within the brain, as demonstrated in various animal models and human studies.^18,19^ The capability of LIFU to precisely target circuits, such as the PLC to NAc core circuit, presents a significant advancement in neuromodulation, offering insights into both normal and pathological brain states.

LIFU’s potential to influence neurotransmitters, especially in the context of circuit-based neuromodulation, has been substantiated in animal models. For example, Xu et al. (2020) found that LIFU targeting the striatum in a Parkinsonian mouse model enhanced dopamine content as measured in this same brain region using High-Performance Liquid Chromatography (HPLC).^20^ Similarly, Min et al. (2011) used microdialysis to demonstrate that LIFU targeting the thalamus significantly increased dopamine concentrations as assessed in its projection region to the striatum.^21^ While encouraging, these studies are not able to capture rapid, real-time fluctuations in neurotransmitter release. Our study bridges this gap by employing fast-scan cyclic voltammetry. This technique allows for the real-time measurement of neurotransmitter dynamics to understand the time course of LIFU to inhibit dopamine.^22^

The goal of this study was to determine the effect of applying LIFU to the PLC on real-time changes in dopamine release in its projection region to the NAc core. We explored three different LIFU spatial peak pulse average intensities, I_sppa_ (6.5 W/cm^2^, 13 W/cm^2^, and 26 W/cm^2^), to determine the conditions most effective in inhibiting dopamine release, as measured using fast-scan cyclic voltammetry (FSCV).^23,24^ We also measured dopamine release in adjacent regions of the striatum (NAc shell, caudate) in order to establish circuit specificity and determined effects on NAc core dopamine release following LIFU modulation of adjacent cortical and sub-cortical regions (i.e., caudate and somatosensory cortex). We hypothesized that higher LIFU intensities applied to the PLC would result in greater suppression of dopamine release within the connected NAc core circuit. We further hypothesized that effects on dopamine release would be specific to LIFU targeting the PLC, with observable changes in the NAc core but not in the NAc shell. Given the role of the PFC-NAc dopaminergic circuit in reward-related neuropsychiatric disorders, elucidating the intensity-dependent and circuit-specific effects of LIFU on dopamine inhibition holds promise for diverse therapeutic applications.

## Methods and Materials

### Chemicals and Materials

Sodium chloride, sodium sulfate, calcium chloride, sodium phosphate monohydrate, and paraformaldehyde were purchased from Sigma-Aldrich (St. Louis, MO). Potassium chloride and magnesium chloride were purchased from Thermo Fisher Scientific (Waltham, WA). Dopamine hydrochloride was dissolved in 0.1 M HClO_4_ to make a 10 mM stock solution. Phosphate buffered saline (PBS; (131.25 mM NaCl, 3.00 mM KCl, 10 mM NaH_2_PO_4_, 1.2 mM MgCl_2_, 2.0 mM Na_2_SO_4_, and 1.2 mM CaCl_2_) was used at pH 7.4 to dilute the dopamine stock solution to 1.0 μM. De-ionized water (EMD Millipore, Billerica, MA) was used to prepare all aqueous solutions.

### Construction of Carbon Fiber Microelectrodes

Carbon fiber microelectrodes were made by aspirating single 7 μm T-650 carbon fibers (Cytec, Woodland Park, NJ) into glass capillaries (A&M Systems, Inc., Carlsborg, WA) and pulled into two using a vertical PE-22 Electrode Puller (Narishige, Tokyo, Japan). Under a microscope, the extended fibers were manually trimmed to 150–200 μm from the glass seal using a scalpel. Before each use, electrode tips were soaked in 70% v/w isopropyl alcohol for 24 hours and backfilled with 1 M KCl.

### Overall Experimental Design

All animal experiments were performed as approved by the Animal Care and Use Committee (ACUC) of the University of Virginia. A total of 38 male Sprague Dawley rats (Charles River Laboratories, Wilmington, MA, USA) between 280 - 320 g were used across a total of seven experimental conditions. Rats were anesthetized with urethane (0.3 ml/l00 g, 5% saline solution., i.p.) prior to each experiment. The rectal and core body temperature was maintained at 32°C using an isothermal pad (Delta Phase Pad; Braintree Scientific, Braintree, MA, USA). Hourly checks (paw pinch) were made on respiration and depth of anesthesia. During surgery, a local anesthetic (bupivacaine) was used on exposed skin and muscle tissue. After each experiment, animals were euthanized using a guillotine (World Precision Instruments, Sarasota, Florida, USA).

In all experiments, anesthetized animals were fixed with stereotaxically navigated electrodes for fast-scan cyclic voltammetry measurements and a LIFU transducer via holes drilled through the skull. The first condition (n=5) was an inactive sham condition where no LIFU was delivered while recording dopamine from the NAc core. The second (n=7), third (n=7), and fourth (n=7) conditions were active, where LIFU was delivered to the PLC at three different intensities while recording dopamine levels from the NAc core (see LIFU Transducer and Waveform below for details on LIFU stimulation). The fifth condition (n=4) was the first anatomical control region, where LIFU was delivered to the somatosensory cortex (S1J) while recording dopamine levels from the NAc core, which are not expected to be functionally connected. The sixth condition (n=4) was the second anatomical control where LIFU was delivered to the PLC, but dopamine was recorded from the caudate putamen (CP), which again are not expected to be functionally connected. The seventh condition (n=4) and final anatomical control consisted of LIFU being delivered to the PLC while dopamine was recorded from the NAc shell, not the NAc core.

### LIFU Transducer and Waveform

A 10 MHz miniature-case immersion transducer (XMS-310-B, Olympus-ims, Waltham, MA) with a 2 mm element diameter and 6 mm focal length from the exit plane was used for all experiments. Ultrasonic waveforms were generated using a two-channel, 30 MHz Dual Channel Function/Arbitrary Waveform Generator (4054B; BK Precision, Yorba Linda, CA, USA). Channel 1 was a 5-volt peak-to-peak (Vpp) square wave of 1 kilohertz (kHz) (N=500) with a 36% pulse duty cycle (PDC). This setup was used to gate channel 2, which was a 10 MHz sine wave. The overall setup resulted in a 500 ms pulse train of 500 pulses or a pulse repetition frequency (PRF) of 1 kHz, each pulse occurring for 360 microseconds. The output of channel 2 was sent through a 30-watt amplifier (E&I 230L; Electronics and Innovation, Rochester, NY, USA) before being sent to the transducer through its corresponding matching network. Waveforms were visualized using a 200MHz 4 Channel 2GS/s Oscilloscope (TDS2024C; Tektronix, Beaverton, O, USA). These pulsing parameters have been shown previously to induce local suppression of neuronal activity in different human brain areas.^25–27^ LIFU delivery was performed in over a 120-second total sonication duration with 500 ms pulse trains separated by a 4.5-second pulse train interval for a total of 24 pulses, leading to a pulse train repetition frequency (PtRF) of 0.1 Hz and a pulse train duty cycle (PtDC) of 3.6%. The baseline spatial peak pulsed average intensity (I_sppa_) delivered to the target was 13 W/cm^2^, equating to 0.71 megapascals (MPa) and a mechanical index (MI) of 0.44. Two other intensities were employed in separate conditions to evaluate the effect of halving (I_sppa_ = 6.5 W/cm^2^, 0.47 MPa, MI = 0.29) or doubling (I_sppa_ = 26 W/cm^2^, 1.06 MPa, MI = 0.65) the spatial peak, pulsed average intensity on dopamine levels. A pictographic representation of the parameters can be seen in Supplementary Figure 1. Thermal modeling of the temperature rise was used to ensure thermal safety (see Ultrasound Thermal Modelling below).

### Electrode Placement and LIFU Targeting

The rat was placed in a stereotaxic frame, and holes were drilled precisely in the skull to place the stimulating electrodes, working electrodes, reference electrodes, and ultrasound transducer, according to the atlas of Paxinos and Watson^28^. Specifically, the carbon-fiber working electrode was lowered into the NAc core (+1.3 mm anterior-posterior [AP], + 2.0 mm medial-lateral [ML], − 7.1 mm dorsal-ventral [DV]), and the bipolar stimulating electrode (Plastics One, Roanoke, VA, USA) was lowered to the VTA (−4.7 mm AP, + 0.9 mm ML, − 8.5 mm DV). The dorsoventral coordinate of the electrodes was adjusted for maximally stimulated release. A 3.0 mm wide hole was drilled through the skull for the transducer placement to sonicate the dmPFC (+3.6 mm AP, + 0.6 mm ML) for the inactive sham (No LIFU) and each of three real LIFU conditions (6.5 W/cm^2^, 13 W/cm^2^, 26 W/cm^2^). In the first anatomical control, the hole for the transducer was drilled at the primary somatosensory cortex (3.2 mm AP, + 0.6 mm ML), and dopamine was measured in the NAc core. In the second and third anatomical controls, dopamine was measured at the Caudate-Putamen (1.3 mm AP, 2.0 mm ML, − 4.5 mm DV) and NAc shell (1.3 mm AP, 1.0 mm ML, − 6.5 mm DV) respectively, while LIFU was applied to the PLC. An Ag/AgCl wire reference electrode was inserted on the contralateral side of the brain. To electrically stimulate dopamine, a constant biphasic current stimulus at +300 μA, 2 ms, 24 pulses were delivered to the VTA by a bipolar stimulating electrode (Plastics One, Inc., Roanoke, VA, USA). Dopamine was detected using FSCV, stimulations were delivered every 5 minutes, and dopamine release was measured at the NAc core (or control region) for 2.5 hours.

### Fast-scan cyclic voltammetry and Electrochemical measurements

Cyclic voltammograms of dopamine were obtained using a Chem-Clamp potentiostat (Dagan Corp., Minneapolis, MN) coupled to a UNC breakout box (UNC Electronics Shop, Chapel Hill, NC) with a 1 MΩ head stage. HDCV software (UNC at Chapel Hill) was used for data acquisition and analysis. Ag/AgCl wires were used as the reference electrodes. A 3-KHz low-pass filter was used to measure dopamine with a waveform that scanned from −0.4 to 1.3 V and back at 400 V/s at 10 Hz. Measurements were made in 5-minute increments from 30 minutes before LIFU to 120 minutes post-LIFU. LIFU was delivered at time zero, and the first post-LIFU recording was at 5 minutes. Carbon Fiber Micro Electrodes were post-calibrated with 1.0 µM dopamine in PBS following animal experiments.

### Empirical Acoustic Field Mapping

The acoustic pressure field of the transducer was quantified in an acoustic test tank filled with de-ionized, degassed, and filtered water (Ultrasonic Measurement System V3 (UMS3) Precision Acoustics Ltd Dorchester, UK) with a calibrated hydrophone (HN - 0500, Onda Corp., CA, USA). The hydrophone was mounted on a motorized stage and used to measure the pressure from the LIFU transducer in the acoustic test tank. The transducer was positioned in the tank using a custom setup and leveled to ensure the hydrophone was perpendicular to the surface of the transducer. XY and XZ planar scans were performed at an isotropic 0.1 mm resolution. A voltage sweep from input voltages of 20-500 millivolts peak-to-peak (mVpp) in 10 mVpp increments was performed to determine the necessary input voltage to obtain the desired intensity.

### Ultrasound Thermal Modelling

Due to the high fundamental frequency used (10 MHz), there is the potential for ultrasound to create unwanted heating. To predict heating in tissue, we simulated heat transfer using the Pennes bioheat equation.^29^ We ran a one-dimensional model assuming constant properties for uniform brain tissue temperature distribution and solved the equation using a finite difference method. Temperature rise was modeled using the exact parameters tested, including 2 minutes of 10 MHz ultrasound at three different intensities (6.5 W/cm^2^, 13 W/cm^2^, and 26 W/cm^2^), using a 500-millisecond pulse train at a pulse train interval of 4.5 seconds.

### Staining and imaging for tissue damage

After the dopamine measurements, rats (n=2) from the no-LIFU, 6.5 W/cm^2^, 13 W/cm^2^, and 26 W/cm^2^ LIFU groups were perfused with PBS, followed by 4% paraformaldehyde. Brains were then collected and fixed in 4% PFA for 24 hours. 1 mm thick sections of the PLC and the NAc core were then taken using a brain matrix and dehydrated in 70% ethanol for sectioning. After being embedded into paraffin at 58−60°C, the tissue specimen was sliced using an automatic microtome and then stained using Hematoxylin and eosin (H&E) at the University of Virginia histology core. The prepared histological sections were observed with an optical microscope (ZEISS Axio Examiner 2), and the images were analyzed using the Image J nuclei counter plug-in.

## Results

### Ultrasound Characterization

Fig. 1A shows a pseudo color axial (Z) pressure map overlaid to scale on a rat brain atlas to show overlap with the desired sonicated area (PLC). The lateral XY and XZ pressure maps are shown in Figure 1B and 1C. The full-width half maximum (FWHM) in the XY plane was ∼ 0.7 mm, while the FWHM in the Z axis was 6 mm. Ultrasound heating models were run using three different intensities. Supplementary Figure 2 shows the predicted temperature rise over the 2-minute sonication. The starting temperature was set at 37°C. The maximum predicted temperature rise for 6.5 W/cm^2^ was 37.2°C, 13 W/cm^2^ was to 37.4°C, and 26 W/cm^2^ to 37.9°C. These temperature rises would not constitute a thermal mechanism and are less than the natural brain fluctuation.^30^

**Figure 1:**
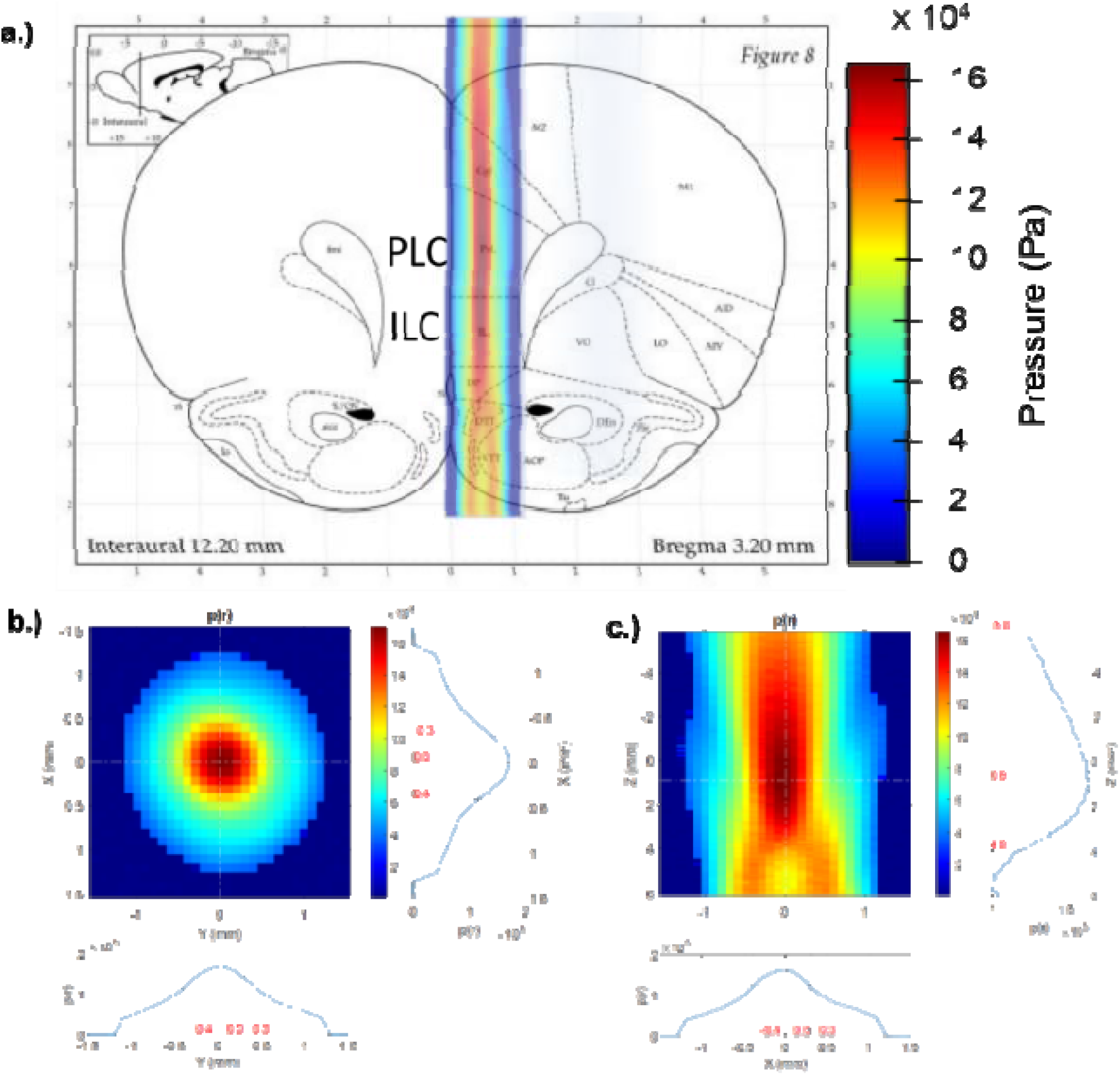
Characteristics of Low-Intensity Focused Ultrasound Sonication. **a).** Overlay of coronal rat brain section and LIFU acoustic waves z-plane showing highest LIFU intensity at pre-limbic region. **b.)** x-y planes of LIFU acoustic waves showing peak intensity focused within a ∼0.7 mm diameter. Vertical dashed lines online plots show a -3dB range. **c.)** x-z planes of LIFU acoustic waves showing peak intensity focused at ∼6-7 mm from the exit plane of the transducer (bottom pointing up). Vertical dashed lines online plots denote the -3dB range.

### Stimulated Dopamine Control

Figure 2A shows the experimental setup for the no-LIFU control experiments where the carbon-fiber recording electrode is placed at the NAc core, and the stimulating electrode is placed at the VTA. Figure 2B shows typical stimulated dopamine release with a color plot, concentration vs. time trace, and cyclic voltammogram.^24^ Dopamine rises when stimulated and decreases to baseline in a few seconds. The color plot and voltammogram identify the species detected as dopamine.^23^ Dopamine release was stable over the first hour and slightly decreased after that, decreasing to 83 ± 2% compared to the first 30 minutes baseline (Fig. 2C). Following the calibration of the electrodes after the experiment, an average dopamine release of 200 ± 9 nM (range: 170 nM – 313 nM) was observed.

**Figure 2:**
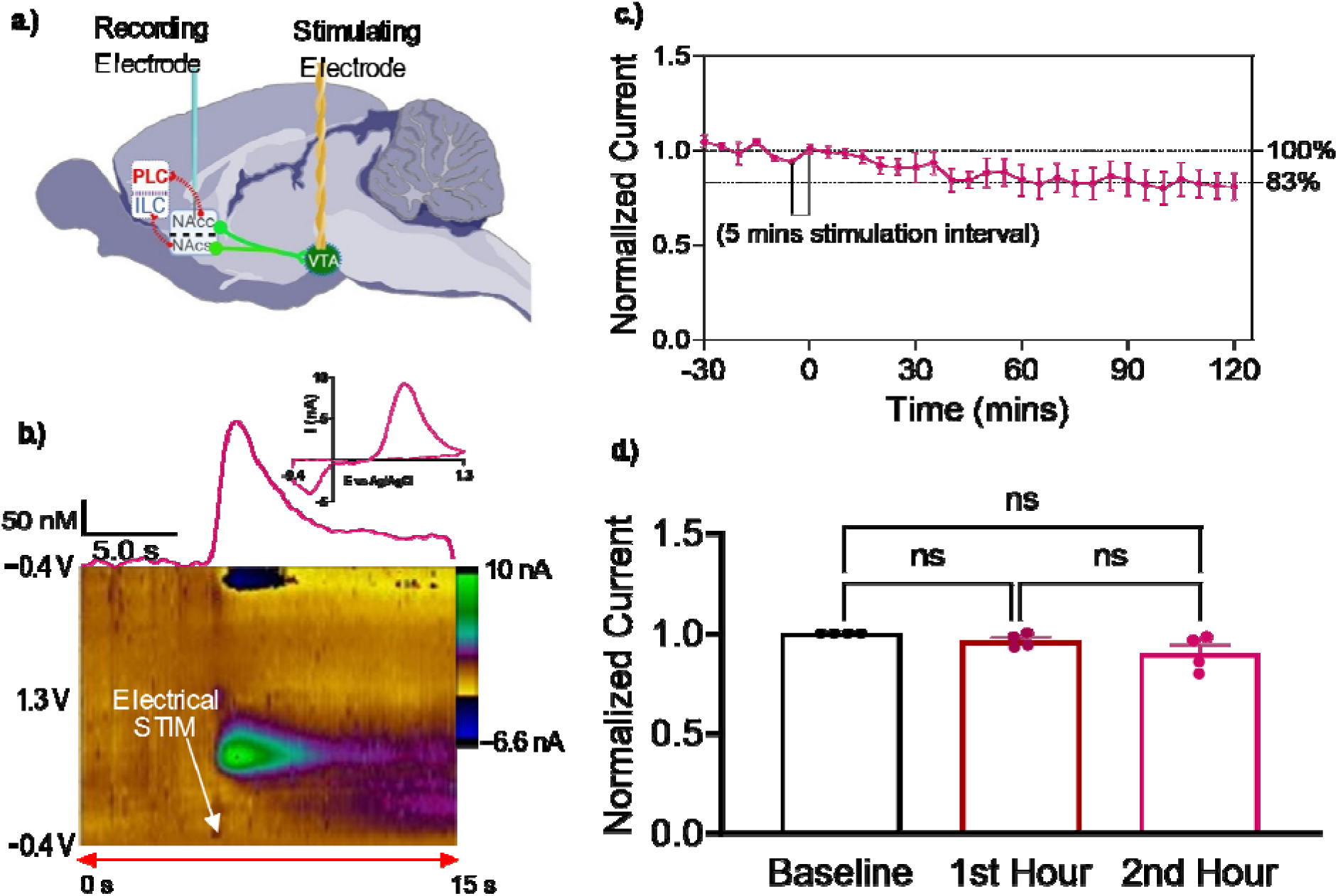
Stimulated Dopamine Control. **a)** Experimental setup: For the no-LIFU control experiments, the carbon-fiber recording electrode was placed at the NAc core, and the stimulating electrode was placed at the VTA. **b) (Top)** Concentration vs. time curve. The color plot and voltammogram show 200 ± 4 nM dopamine release at the carbon-fiber microelectrode (recording electrode) surface using FSCV. **c)** Group (n = 4) mean ± SEM dopamine release over time was measured every 5 minutes. After 2 hours, the dopamine release reduction was 17 ± 0.3 %. **d)** Group (n = 4), mean ± SEM, data from C grouped into baseline and 1-hour blocks. Dopamine release did not significantly change in the first hour and second-hour post-LIFU sonication as compared to baseline.

### PLC LIFU Sonication Causes Dopamine Release Inhibition in the NAc Core

Fig. 3B shows the dopamine data with 13 W/cm^2^ intensity, including the color plot concentration vs. time curves (Figure 3B). The color plot shows the dopamine release is lower (less green color intensity), and the concentration vs. time curves show the peak current decreased from 7 nA to 3.5 nA. The dopamine concentration dropped from 160 ± 10 nM to 82 ± 5 nM, a 50 ± 3 % decrease. Figure 3C shows the averaged data for n = 7 rats, and inhibition was observed 2 minutes after sonication with 13 W/cm^2^ LIFU, and dopamine release continued to be inhibited for 2 hours after a single LIFU treatment. Fig. 3D shows that LIFU sonication of the PLC evoked a 42 ± 3 % dopamine inhibition in the first hour and an additional 7 ± 1 % in the second-hour post-LIFU sonication.

**Figure 3:**
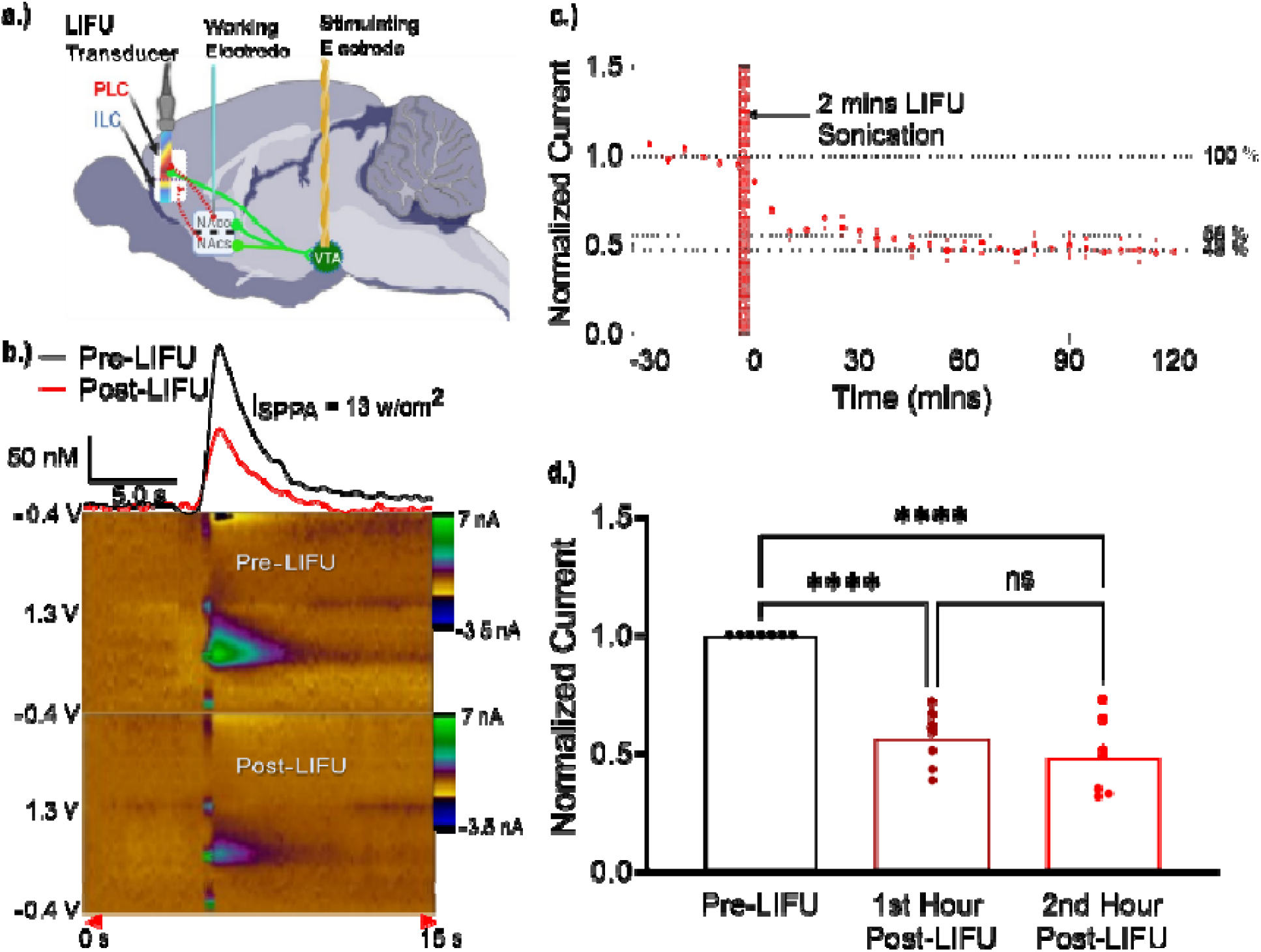
Prelimbic Region LIFU Inhibited Dopamine Release in the Nucleus Accumbens. **a)** The LIFU transducer was placed above the brain for the PLC sonication at 13 W/cm^2^. **b)** Color plots and concentration-time trace show the inhibition of dopamine release from 160 ± 10 nM to 82 ± 5 nM after 2 hours post-LIFU. **c)** Group (n = 7) means ± SEM. Normalized current over time. The vertical red bar denotes the application of 2 minutes of LIFU to PLC. Dopamine release was tested every 5 minutes out to 2 hours after LIFU. **d)** Group (n = 7) mean ± SEM. Data from C collapsed into baseline and 1-hour bins. Focused ultrasound sonication of the PLC invoked a 42 ± 3 % average dopamine inhibition in the first-hour post-LIFU sonication that did not significantly change (7 ± 1 %) in the second hour.

### Effect of LIFU Intensity Dosing on Dopamine Inhibition

To investigate the effect of LIFU intensity on dopamine release, we halved (6.5 W/cm^2^) and doubled (26 W/cm^2^) the LIFU intensity while leaving all other parameters constant. Figure 4A shows that the experimental design used is the same as the setup in Figure 3A. Figure 4B shows the overlay of the concentration vs. time traces for dopamine release before and after LIFU sonication for both spatial peak average intensities. Dopamine release dropped from 180 ± 7 nM at baseline to 144 ± 5 nM two-hour post-LIFU at the 6.5 W/cm^2.^ intensity and from 150 ± 5 nM at baseline to 100 ± 3 nM two-hour post-LIFU at the 26 W/cm^2^ intensity (Figure 4A). Fig. 4B shows that 6.5 W/cm^2^ *I_SSPA_* resulted in only about 20 ± 1% reduction in dopamine release, unlike the immediate drop in stimulated dopamine release with 13 W/cm^2^, similar to the no-LIFU controls. Applying 26 W/cm^2^ *I_SSPA_* resulted in a 30 ± 1% gradual inhibition, taking about 45 minutes to reach the point of maximal decrease (Figure 4C). In Figure 4D, sonicating the PLC with 13 W/cm^2^ *I_SSPA_*had the highest inhibitory effect on dopamine release in the NAc core. 6.5 W/cm^2^ *I_SSPA_*showed no significant dopamine release inhibition 2 hours post-sonication.

**Figure 4:**
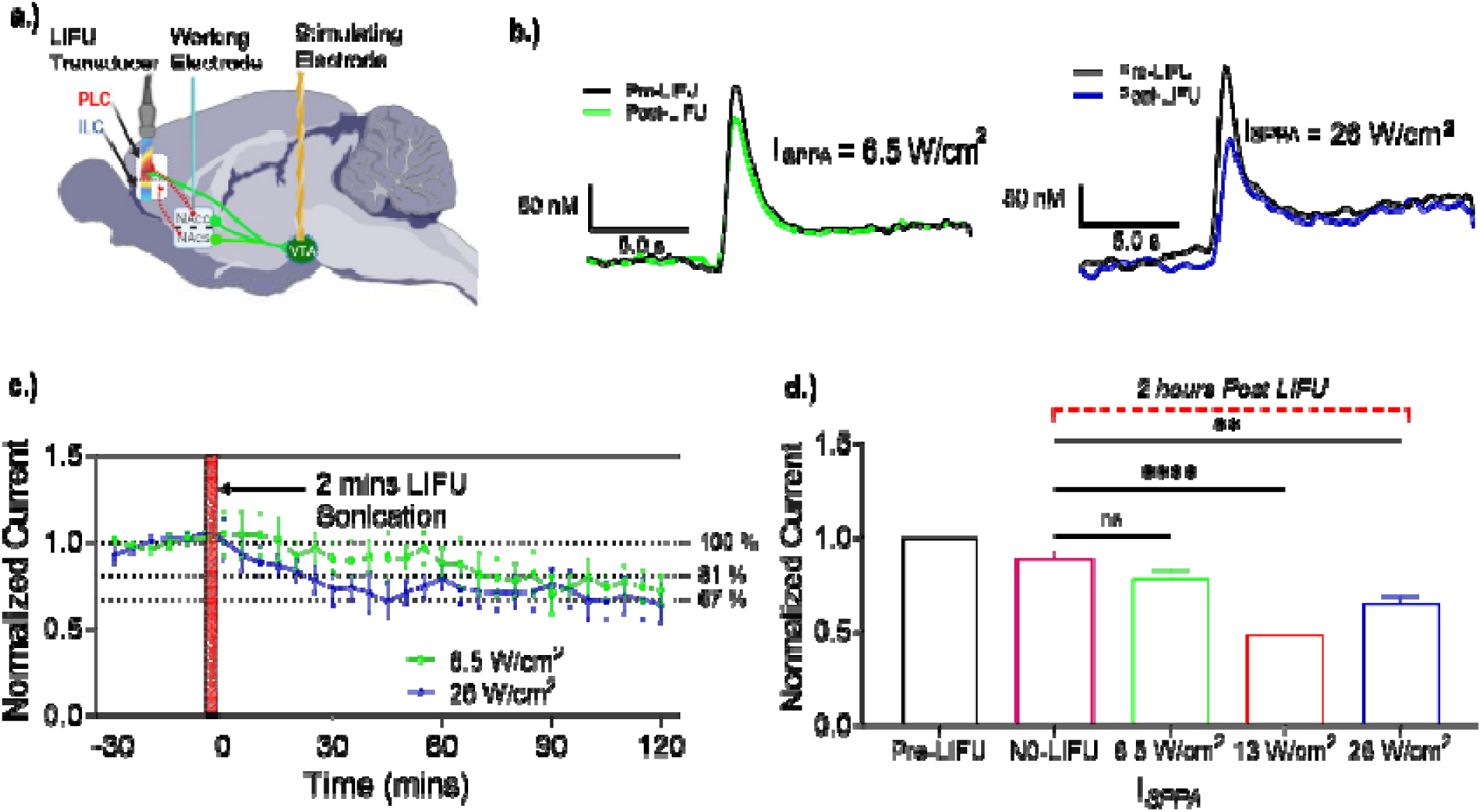
Intensity-Dependent Effects of LIFU on Dopamine Release in the Nucleus Accumbens. **a.)** The LIFU transducer was placed above the brain for the PLC sonication at both 6.5 and 26 W/cm^2^ **b.)** Concentration vs. time traces overlay and time-concentration scale bar show the change of dopamine release upon LIFU sonication. Sonication of PLC with 26 W/cm^2^ I_SPPA_ reduced the dopamine release from 150 ± 5 nM to 100 ± 3 nM. The average dopamine release reduced from 180 ± 7 nM to 144 ± 5 nM with 6.5 W/cm^2^ **c.)** As a confirmation of the overlayed I-t curves, the 2-hour-long dopamine stimulation shows a 30 ± 1.0% change with 26 W/cm^2^ I_SPPA_ and a 20 ± 0.7 % change with 6.5 W/cm^2^ I_SPPA,_ both with a 5-minute stimulation interval. **d.)** Bar graph of a summarized result of dopamine changes 2 hours post-LIFU sonication with the tested LIFU parameters—in the order of dopamine inhibition, 13 W/cm^2^ I_SPPA_ (n=7) > 26 W/cm^2^ I_SPPA_ (n=7) > 6.5 W/cm^2^ I_SPPA_ (n=7, separate groups for each parameter).

### Circuit Specificity for the Inhibitory Effects of LIFU

To ensure that LIFU precisely inhibits the dmPFC circuit, specifically as opposed to the potential for non-specific effects,^31^ we conducted three anatomical control experiments, as shown in Figure 5A. In all cases, the brain was sonicated with 13 W/cm^2^ *I_SSPA_* since it gave the highest inhibition of dopamine among the tested intensities. Since S1J does not directly regulate NAc core dopamine release,^32^ and the PLC does not regulate dopamine release in the CP or NAc shell,^33^ we expected no significant inhibition of dopamine release in all cases. As expected, dopamine release did not notably decrease in the S1J-NAc core, as it decreased from 150 ± 45 nM to 127.5 ± 4 nM, which is similar to controls (Fig. 5B). For the PLC-CP and the PLC-NAc shell control, dopamine release decreased from 160 ± 3 nM to 138 ± 3 nM and 118 ± 4 nM to 101 ± 3 nM, respectively, also similar to controls (Fig. 5B). On average, dopamine levels reduced gradually by 25 ± 1 % in the S1J-NAc core control, 14 ± 1 % in the PLC-CP control and 19 ± 1 % in the PLC-NAc shell control in 2 hours post-sonication (Figure 5C). Figure 5D shows that there is no significant difference between the dopamine level reduction with the No LIFU stimulated dopamine control and each anatomical control.

**Figure 5:**
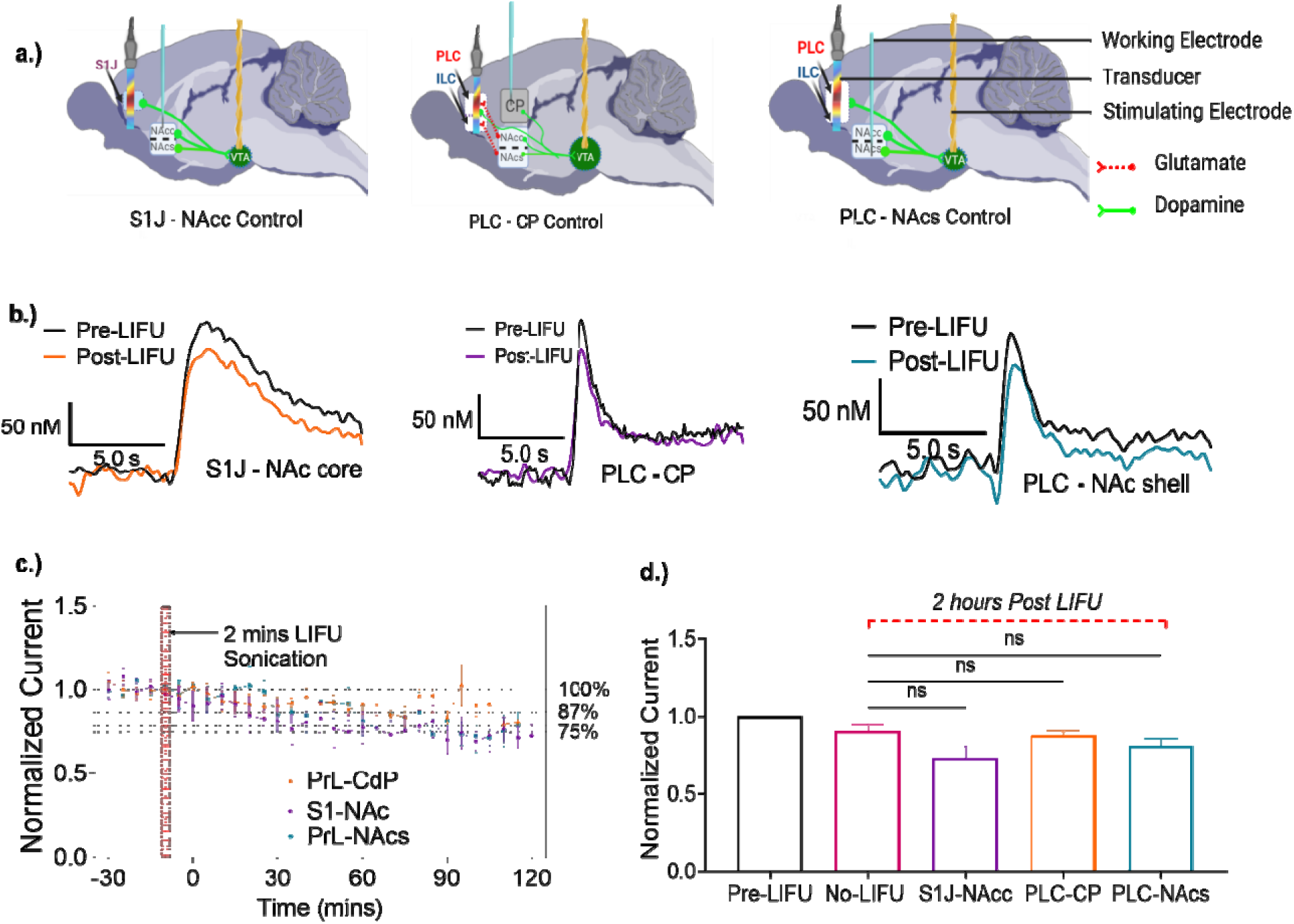
Circuit-Specificity of LIFU Neuromodulation. **a.)** Experimental Setup for control experiment **b)** Concentration vs. time traces for anatomical controls for LIFU sonication with 13 W/cm^2^ I_SPPA_. Top: S1J-NAc core control showed a dopamine concentration change of 150 ± 5 nM to 128± 4 nM. Middle: PLC-CP control showed a concentration change of 160 ± 3 nM to 138 ± 3 nM. Right: PLC-NAc shell control showed a concentration change of 118 ± 4 nM to 101 ± 3 nM **c)** Dopamine stimulation data shows a 25 ± 0.8 % change in the S1J-NAc core control, a 14 ± 1 % change in the PLC-CP control, and a 19 ± 1 % change in the PLC-NAc shell control with 13 W/cm^2^ I_SPPA_ after 2 hours. **d)** Bar graph of a summarized result of dopamine release change after 2 hours post-LIFU sonication in the anatomical controls (n = 4 animals per treatment) compared to the No-LIFU control (n = 4). There was no significant difference between the first and last dopamine levels after the 2 hours of LIFU sonication.

### Statistical Modelling of Intensity-Dependency and Circuit Specificity of LIFU on Dopamine Inhibition

A generalized linear mixed-effects model was applied using MATLAB. This model was chosen to include all predictor variables (treatment/control condition, time and time-condition interaction). Statistical modeling of the data showed significant differences from the null (F _(13,1126)_ = 175.98, p < 0.0001) with an adjusted R^2^ (R^2^_adj_) = 0.67. A significant main effect of Time (F _(1,1126)_ = 37.05, p < 0.0001) and the Condition*Time interaction (F (1,1126) = 13.46, p < 0.0001) was observed, with no main effect of Condition (F _(1,1126)_ = 1.1019, p = 0.411) (Supplementary Table 1). Breaking down the interactions demonstrated two LIFU conditions (medium, and high intensities) significantly decreased dopamine levels over time as compared to the reference (No LIFU) condition. A breakdown of the beta coefficients, standard errors, t-statistics, and p-values for all effects is shown in Supplementary Table 2.

A plot of all 7 conditions across time can be seen in Figure 6A. After correction for multiple comparisons (threshold of p < 0.0167), linear hypothesis testing of the significant interactions revealed that the slopes of the interaction of the medium intensity with time (13 W/cm^2^:Time) and high intensity with time (26 W/cm^2^:Time), but not low intensity with time (6.5 W/cm^2^:Time) showed significantly decreased dopamine levels over time compared to the reference (No LIFU) condition (Figure 6B).

**Figure 6.**
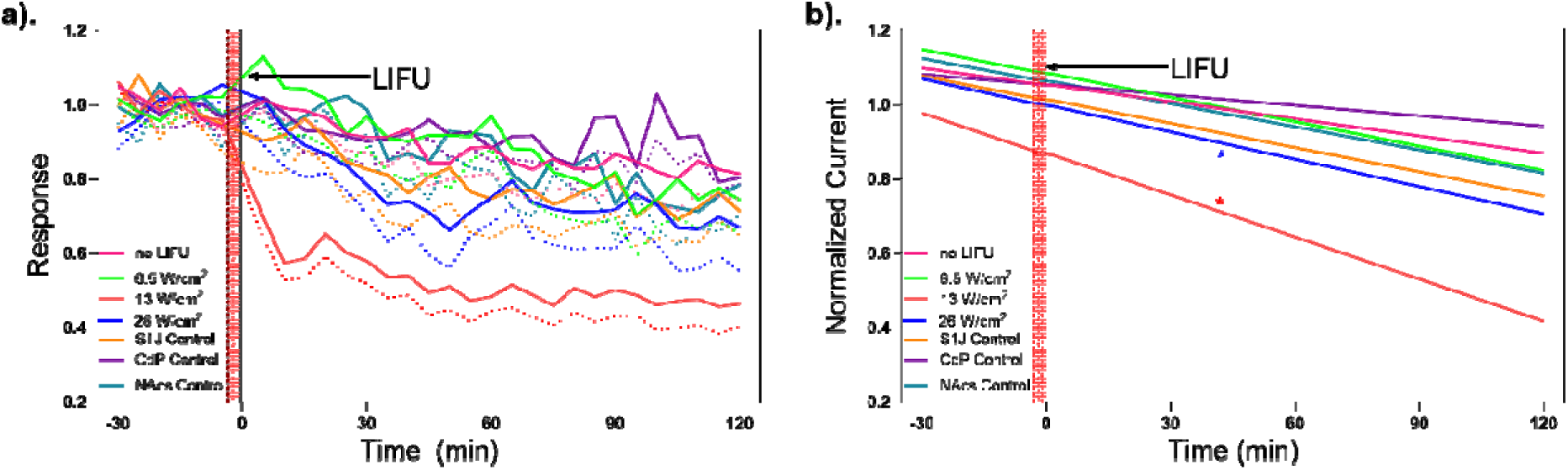
Dopamine levels over time and modeling results. **a.)** Group means ± SEM normalized dopamine levels across time for each of the 7 experimental conditions. The Y-axis is normalized dopamine levels; the X-axis is time in minutes (min). Time point zero represents the LIFU application. The solid black line represents the no low-intensity focused ultrasound (No LIFU) or inactive sham condition. Solid red, green, and blue lines represent the low-intensity (6.5 W/cm2), medium-intensity (13 W/cm2), and high-intensity (26 W/cm^2^) LIFU conditions. Dashed yellow, purple, and orange lines represent the somatosensory (S1J), caudate-putamen (CdP), and nucleus accumbens shell (NAcs) active control conditions. The vertical orange bar represents the LIFU application period. **b.)** Modelled slopes of each condition for the post hoc linear hypothesis testing. Colored stars represent the conditions significantly different from the reference (No LIFU) condition after correction for multiple comparisons.

### No Effect of LIFU on Markers of Cell Damage or Death

Figure 7 shows standard bright field images of Hematoxylin and Eosin (H&E) -stained coronal sections of the PLC. As shown by the intactness of most nuclei (purple), no significant cell damage was observed with the No-LIFU Control (Figure 7A), 6.5 W/cm^2^ LIFU *I_SSPA_* (Figure 10B), 13 W/cm^2^ LIFU *I_SSPA_* (Figure 7C), or 26 W/cm^2^ LIFU *I_SSPA_* (Figure 7D). The bar graph in Figure 7E inset compares the nuclei count of the right brain PLC (LIFU-sonicated) to the left-brain PLC of each animal (control). Mixed-effects analysis in Figure 7E shows no significant difference between the cell count of the treated (right) and untreated hemispheres (left) of any groups (F _(4,3)_ = 4.76, p=0.9412, No-LIFU control, p=0.9893 for 6.5 W/cm^2^ I_SPPA_, p>0.9999 for 13 W/cm^2^ I_SPPA_, and p=0.2557 for 26 W/cm^2^ I_SPPA_). Figure 7F compares all the nuclei counts between the nuclei counts of the sonicated PLC versus the No-LIFU controls. There was no significant difference in the nuclei counts of the sonicated PLC 1-hour post-test compared to control (One-Way ANOVA; F _(1.5, 4.6)_ = 3.945, p = 0.3959 for 6.5 W/cm^2^ I_SPPA_, p = 0.1701 for 13 W/cm^2^ I_SPPA_, and p = 0.6366 for 26 W/cm^2^ I_SPPA_).

**Figure 7.**
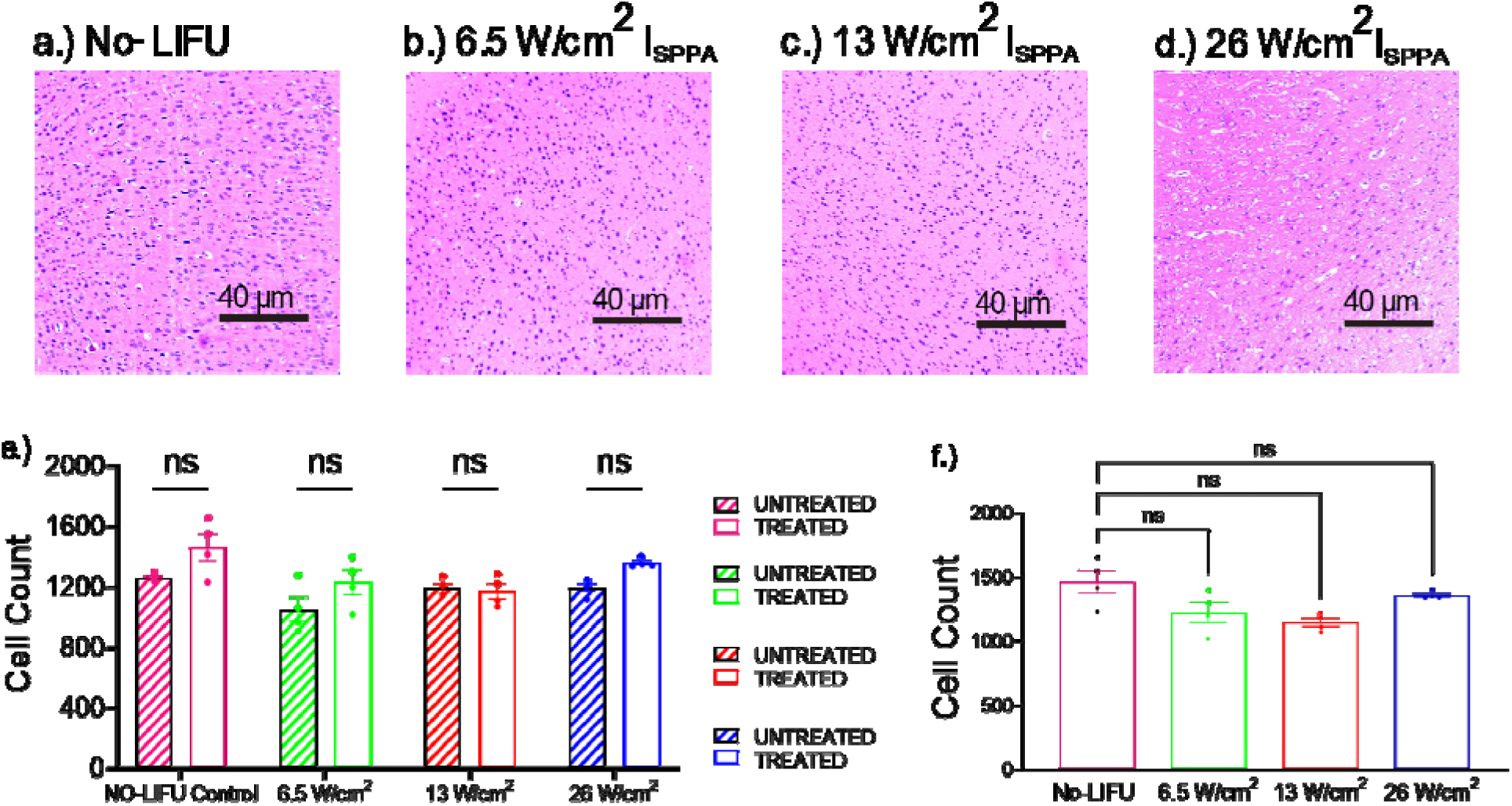
LIFU does not result in cell damage or death. Comparison of LIFU sonicated right pre-limbic area with non-sonicated left pre-limbic area. **a.)** No-LIFU Control. **b.)** 6.5 W/cm^2^ LIFU I_SSPA_. **c.)** 13 W/cm^2^ LIFU I_SSPA_. **d.)** 26 W/cm^2^ LIFU I_SSPA._ **e.)** The bar graph shows no significant change in the treated (right) brain pre-limbic region LIFU sonicated nuclei count versus the left (untreated) brain control pre-limbic region. f.) The bar graph shows no significant change between the nuclei counts of the right brain pre-limbic region of the sonicated brains versus the No-LIFU controls. (n=4 for all conditions).

## Discussion

The purpose of this study was to demonstrate the use of LIFU as a tool for inducing circuit-specific inhibitory neuromodulation of dopamine. We investigated the impact of low-intensity focused ultrasound (LIFU) on real-time dopamine release within the PLC and its projection region, the NAc core. The primary finding was that a 2-minute LIFU treatment of the PLC with an *I_SSPA_* of 13 W/cm^2^ robustly (∼50%) and persistently (2-hr) inhibited dopamine release in the NAc core. LIFU-induced inhibition was intensity-dependent, but not in a linear manner, as doubling the intensity to 26 W/cm^2^ decreased the inhibition to 30%. Reducing intensity to 6.5 W/cm^2^ did not significantly affect dopamine release as compared to controls. The effects were also circuit-specific: LIFU to the somatosensory cortex did not impact dopamine release in the NAc core, and LIFU to the PLC did not affect dopamine release in the caudate or NAc shell. LIFU did not result in appreciable heating and caused no cell damage or death, suggesting it can safely modulate the PLC, resulting in robust downstream neural circuit effects. These findings have strong translational implications, especially considering that ultrasound technology is already FDA-approved for human use. This study, therefore, positions LIFU as a noninvasive, precisely targeted, and effective tool for modulating dopaminergic neural circuits.

### Circuit-Specific Dopamine Inhibition in the NAc Core Induced by PLC LIFU Sonication

This study demonstrates that phasic dopamine release in the NAc core is inhibited by targeting the PLC. A 2-minute application of LIFU sonication to the PLC resulted in persistent suppression of electrically stimulated phasic dopamine release. The most exciting finding from this study was that LIFU modulation is very circuit specific. LIFU applied to the PLC did not change dopamine in the adjacent striatum or NAc shell, while LIFU applied to a region adjacent to the PLC did not change dopamine in the NAc core. While the focused ultrasound beam is centered in the PLC, about 40% of the maximal LIFU intensity may have targeted the infralimbic cortex (ILC). The ILC projects directly to the NAc shell and does not directly affect dopamine release in the core.^33^ As a control, we applied LIFU to the PLC (which could result in 40% LIFU intensity reaching the ILC) and found that it did not affect stimulated dopamine release in the NAc shell. Thus, LIFU effects were restricted to the beam maximum, and the LIFU was applied circuit-specifically.

In the control group without LIFU, we observed a modest 14 -17% decrease in dopamine after 2 hours of VTA electrical stimulation, indicating stable release without LIFU. It is important to note that while the urethane anesthesia used in this study has been reported to impact dopamine release in vitro, urethane does not affect dopamine release or uptake in vivo studies.**^34–37^** Therefore, the observed slight decrease in dopamine release is likely not due to the use of anesthesia. Our experimental design in anesthetized animals limited dopamine measurement to a two-hour timeframe. Nonetheless, the inhibition lasts at least two hours, demonstrating that even a 2-min LIFU treatment leads to persistent neurochemical change.

Many studies have used the NAc core as a read out of activity in the dmPFC.^38–40^ The reduction in dopamine level is proposed to be an effect of the inhibition of PLC-NAc core glutamatergic neurons. This hypothesis is based on previous studies showing that glutamatergic projections from the PLC to the NAc core function in a dopamine-dependent manner, where an increase in the neuronal activity of the PLC gives a corresponding increase in the neuronal activity of the NAc core.^9,41–44^ Thus, our findings potentially represent a downstream consequence of reduced glutamatergic activity from PLC neurons. While we cannot measure glutamate with FSCV because it is not electroactive, we measured the downstream effects on dopamine in the NAc core and saw inhibition.^45–47^ Thus, modulating the PLC with inhibitory LIFU results in downstream inhibition of dopamine release.

### Non-Linear Relationship Between LIFU Intensity and Dopamine Inhibition

LIFU influences parameter-based physiological activity which may determine its mechanisms of action.^18^ While this study was not specifically designed to investigate the mechanism of action, we focused on the effect of intensity, as this parameter may be a driver of LIFU neuromodulation. Lower intensities are typically preferred due to the lower risk of damage, particularly regarding thermal effects, which is a significant concern for humans. To address this, we modeled the temperature rise using a simplified one-dimensional linear model with the Pennes bioheat equation, as previously done in LIFU studies.^48^ The optimal intensity of effect (13 W/cm²) produced a maximum temperature rise of 0.4°C, while the higher intensity (26 W/cm²) produced a maximum temperature rise of 0.9°C (Supplementary Figure 2).

Some literature suggests that LIFU may induce inhibitory responses via thermal mechanisms.^49–54^ For instance, Darrow et al. (2019) demonstrated that temperature rise closely tracked the suppression of somatosensory evoked potentials in rats, with effects becoming more pronounced at a 0.5-1°C rise, similar to our higher-intensity LIFU parameters.^55^ In contrast, Collins et al. (2021) suggested that a 2-3°C temperature rise was needed to produce inhibitory responses in a leech model.^56^ While we cannot rule out a thermal mechanism contributing to LIFU’s observed effects on dopamine levels in this study, higher temperature rises did not produce larger effects on dopamine. Moreover, the optimal intensity produced an estimated heating on the lower end of prior reports on thermal effects. One possible mechanism for this effect is through mechanosensitive ion channels, as LIFU sonication may induce enduring alterations critical for regulating neuronal firing.^57,58^ These alterations may consequently decrease neuronal excitability, leading to a decreased responsiveness to electrical stimulation over a longer period. We propose that the observed effects were likely driven primarily by a mechanical mechanism rather than a thermal one. However, further investigation is warranted to elucidate the precise mechanisms of LIFU-mediated neurotransmission.^59–62^

Our data align with findings by Legon et al. (2018), where the same PRF and duty cycle inhibited evoked potentials in the thalamus at 14.56 W/cm².^63,64^ However, we found a U-shaped effect of LIFU intensity as opposed to other LIFU studies.^65^ The results demonstrate that the medium intensity employed (13 W/cm²) was more effective at reducing dopamine than the highest intensity. Other research indicates that LIFU’s effect follows a sigmoidal relationship with intensity, where the effect plateaus, and additional energy does not increase it.^66,67^ Similarly, a lower inhibition level of dopamine at 26 W/cm² aligns with Hoffman et al. (2022), where evoked action potentials in mouse sensory neurons were reduced between 11 and 30 W/cm² upon activation of the mechanically gated ion channel PIEZO2.^68^ Ion channels do not always exhibit a linear response to neuronal firing due to their complex and contingent nature on several factors.^69^ Nevertheless, these results suggest that higher intensities do not result in greater effects, instead supporting a "sweet spot" where responses are optimally elicited.^70^

This study demonstrates the inhibition of dopamine release by applying LIFU to the mesolimbic circuit in the rat brain. Notably, the use of LIFU to treat substance use disorder by directly modulating the NAc is being explored. For instance, Mahoney et al. (2023) used a 10-minute-long NAc sonication to reduce cue-induced substance craving activity in humans—an indirect indication of local NAc dopamine inhibition.^71^

## Conclusion

In conclusion, we have shown that circuit-specific LIFU-induced dopamine inhibition can be elicited in the mesolimbic pathway with a short duration (two-minute) sonication. This inhibition is intensity-dependent, albeit not linearly. This is the first study to measure real-time downstream LIFU inhibition dopamine release in the mesolimbic pathway, showing the effect of intensity variation and circuit specificity. Thus, our results extend the knowledge of LIFU modulation in treating neuropsychological disorders involving the PLC and NAc. Future studies could explore different LIFU parameters to determine if mesolimbic dopamine could be excited by LIFU. The shorter sonication duration observed in this study could inform parameters for transcranial LIFU in humans. Additionally, these results could be helpful for other disorders involving the NAc and the dopamine system.^32,72,73^

## Statements and Declarations

### Author contribution

Greatness O. Olaitan; Investigation and writing, Mallikarjunarao Ganesana; Equipment Setup and experiment verification, Andrew Strohman; equipment testing, statistics, and manuscript editing Wendy J. Lynch; review-editing, Wynn Legon; equipment testing and manuscript editing, B. Jill Venton; project administration, review-editing.

### Supporting Information

None

### Ethics Approval

All animal experiments were approved by the University of Virginia Institutional Animal Care and Use Committee.

### Source of biological material

All animals were purchased from Charles River.

### Statement on animal welfare

Animals were used in accordance with guidelines on animal welfare published by OLAW from NIH.

