## Supplementary material for "Focused Ultrasound Modulates Dopamine in a Mesolimbic Reward Circuit": Supplemetary File

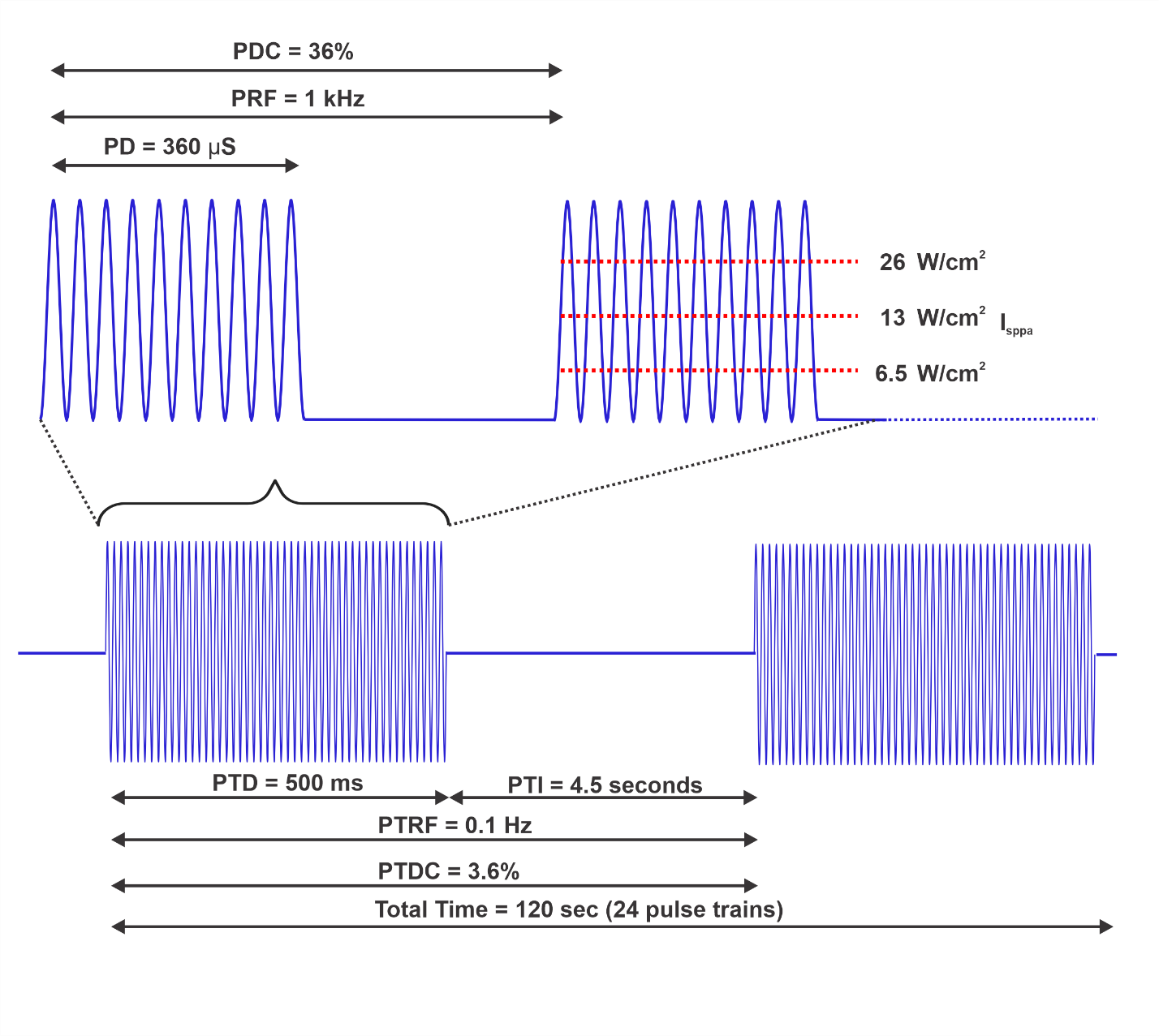
 **Supplementary Figure 1. Low-intensity focused ultrasound (LIFU) Parameters.** **(Top).** Breakdown of each pulse within a single pulse train, including the pulse duration (PD), pulse repetition frequency (PRF), and the pulse duty cycle (PDC). Horizontal dashed red lines represent the spatial peak, pulsed average intensity (I_sppa_) at each of the three active LIFU conditions from low to high intensity in the bracket. **(Bottom).** Breakdown of LIFU application across pulse trains, including the pulse train duration (PTD), pulse train interval (PTI), pulse train repetition frequency (PTRF), the pulse train duty cycle (PTDC), and the total time for LIFU application.

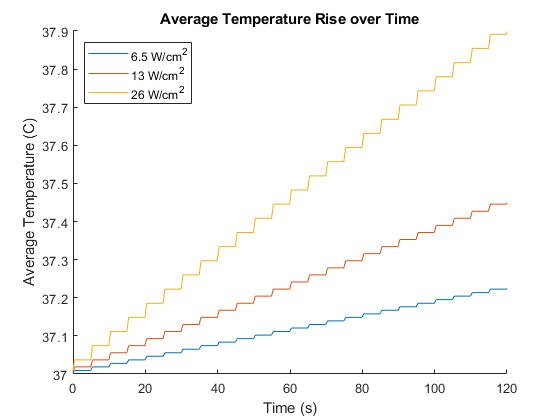

**Supplementary Figure 2: Modeling of ultrasound temperature rise.** The figure shows estimated heating in the brain for the three tested intensities for the 2 minutes of sonication.

***Statistical Analysis***

Raw dopamine concentration values (nM) at each time point were normalized to the mean dopamine levels observed during the 30-minute pre-LIFU baseline period. A generalized linear mixed-effects model was then applied using MATLAB R2022a's fitglme function. The outcome variable was the normalized dopamine levels, while the predictor variables included condition (No LIFU, 6.5 W/cm², 13 W/cm², 26 W/cm², S1J control, CdP control, and NAcs control), time (from 30 minutes before to 120 minutes after LIFU in 5-minute increments), and the interaction between condition and time. The No LIFU (inactive sham) condition served as the reference. To account for individual variability, animals were treated as a random effect. Model assumptions were evaluated using predicted vs. residual plots, Q-Q plots, and residual histogram plots. The model output included the adjusted R-square, F-statistic (degrees of freedom), and p-value for the overall model, as well as F-statistics (degrees of freedom) and p-values for each variable as shown in supplementary table 1.

**Supplementary Table 1. Overall Statistical Model Results**

| **Variable** | **F-statistic** | **DF_num_** | **DF_den_** | **p-value** | **R^2^_adj_** |
| --- | --- | --- | --- | --- | --- |
| Overall | 175.98 | 13 | 1126 | <0.0001 | 0.67 |
| Intercept | 307.676 | 1 | 1126 | <0.0001 |  |
| Condition | 1.019 | 6 | 1126 | 0.411 |  |
| Time | 37.045 | 1 | 1126 | <0.0001 |  |
| Condition: Time | 13.462 | 6 | 1126 | <0.0001 |  |

Additionally, beta coefficients, standard errors, t-statistics (degrees of freedom), and p-values for each fixed effect were reported. The overall model results are summarized in Supplementary Table 1, and the fixed effects are detailed in Supplementary Table 2. Data are plotted as the mean ± SEM. Post hoc analysis of significant results was conducted using a linear hypothesis test (coefTest function) on the beta coefficients of the fixed effects, with Bonferroni correction for multiple comparisons. Data are presented as the slopes for each condition.

Staining and imaging data were analyzed using one-way ANOVA and mixed-effects model analysis in GraphPad Prism v. 10.0 (GraphPad Software, La Jolla, CA). Results are reported as F-statistics (degrees of freedom) and p-values. Values on plots are reported as the mean ± SEM, with 'n' representing the number of animals.

**Supplementary Table 2. Fixed Effects**

| **Fixed Effect** | **Beta** | **SE** | **t-statistic** | **DF** | **p-values** |
| --- | --- | --- | --- | --- | --- |
| Intercept | 1.026 | 0.058 | 17.541 | 1126 | <0.0001 |
| 6.5W/cm^2^ | 0.053 | 0.077 | 0.697 | 1126 | 0.486 |
| 13W/cm^2^ | -0.109 | 0.077 | -1.428 | 1126 | 0.153 |
| 26W/cm^2^ | -0.024 | 0.077 | -0.313 | 1126 | 0.754 |
| S1J Control | -0.018 | 0.088 | -0.206 | 1126 | 0.837 |
| CdP Control | -0.020 | 0.088 | -0.229 | 1126 | 0.819 |
| NAcs Control | 0.029 | 0.088 | 0.328 | 1126 | 0.743 |
| Time | -0.008 | 0.001 | -6.086 | 1126 | **<0.0001** |
| 6.5W/cm^2^: Time | -0.003 | 0.002 | -1.968 | 1126 | **0.05** |
| 13W/cm^2^: Time | -0.011 | 0.002 | -6.718 | 1126 | **<0.0001** |
| 26W/cm^2^: Time | -0.005 | 0.002 | -2.727 | 1126 | **0.006** |
| S1J Control: Time | -0.003 | 0.002 | -1.665 | 1126 | 0.096 |
| CdP Control: Time | 0.003 | 0.002 | 1.591 | 1126 | 0.112 |
| NAcs Control: Time | -0.003 | 0.002 | -1.409 | 1126 | 0.159 |
